# Collagen-Based Tumor Spheroid Model for Investigating Tumor-Macrophage Interactions through Extracellular Matrix Remodeling

**DOI:** 10.1101/2025.10.01.679587

**Authors:** Luna Zhang, Sanab Abdi, Hannah M. Szafraniec, Paolo P. Provenzano, Kathryn L. Schwertfeger, David K. Wood

## Abstract

Macrophages in the tumor microenvironment (TME) can constitute up to 50% of tumor mass and play a critical role in cancer cell proliferation, invasion, and metastasis. While their contribution to extracellular matrix (ECM) degradation through matrix metalloproteinases (MMPs) has been explored, the role of other macrophage-derived factors in ECM remodeling and their impacts beyond degradation remain poorly understood. Here, we describe the development of a 3D collagen-based tumor spheroid model to investigate the impact of peripheral blood mononuclear cell (PBMC)-derived macrophages on cancer cell-ECM and cancer cell-macrophage interactions within the TME. We observed that cancer cells stimulated PBMC-derived macrophages into an M2-like phenotype and that tumor spheroid conditioned macrophages (TSCMs) shifted cancer cell populations toward phenotypes with greater invasion distances and reduced circularity, indicative of increased malignancy. Such observations can be explained by macrophage-mediated ECM remodeling. Specifically, we demonstrate that TSCMs secreted a variety of soluble factors that are known to contribute to ECM remodeling, including ECM degradation and fiber realignment. These processes collectively create a tumor-favoring environment by loosening the collagen matrix and aligning fibers that serve as invasion tracks for migrating tumor cells that facilitate cancer cell migration and invasion. This model provides a robust platform to study the interactions between cellular and non-cellular components in the TME and to identify the molecular mechanisms underlying cancer progression. These insights may aid in the development of novel therapeutic strategies targeting macrophage-mediated processes in cancer.

**Graphical Abstract:** 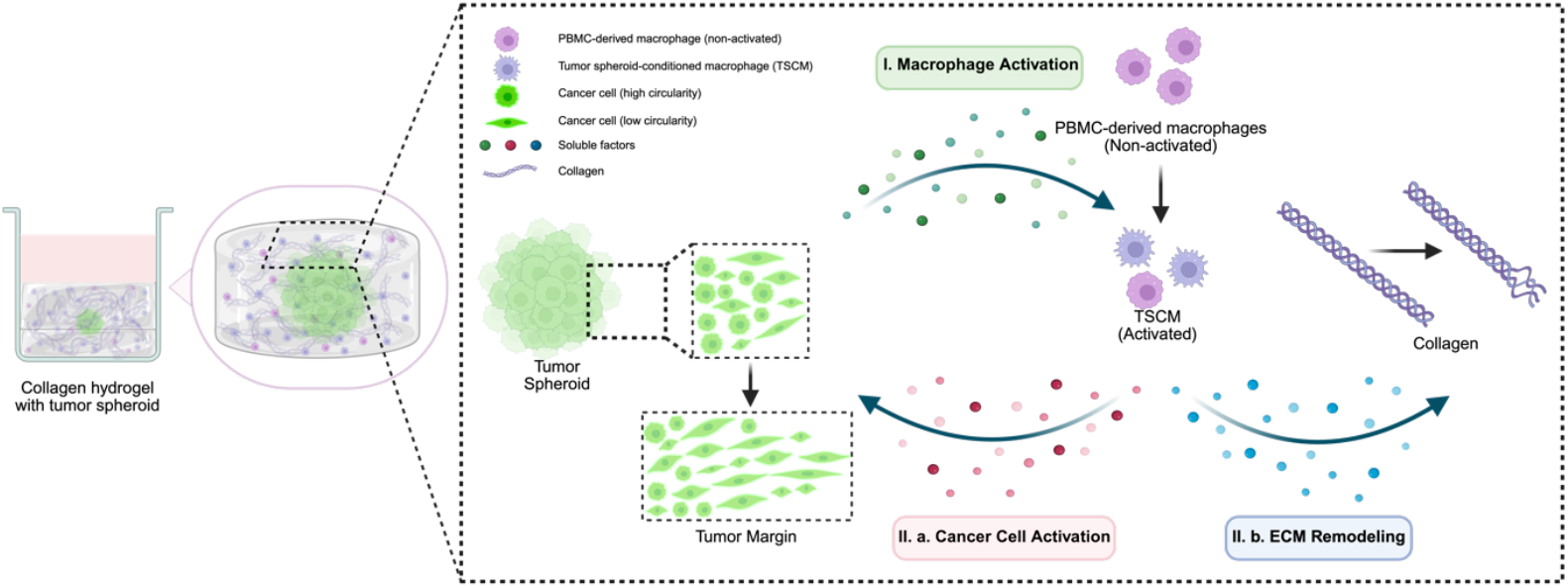

## 1. Introduction

Cancer metastasis, a process that begins with the dissemination of cells from primary tumors, is significantly influenced by the tumor microenvironment (TME). The TME is a highly structured ecosystem composed of cancer cells, stromal cells, extracellular matrix (ECM), and signaling molecules [1]. It provides structural support, mediates biochemical signaling, and dynamically interacts with cancer cells to influence tumor growth, invasion, metastasis, and resistance to therapy [1], [2]. Within this ecosystem, tumor-associated macrophages (TAMs) are crucial players, comprising up to 50% of tumor mass and associated with poor clinical outcomes in breast cancer [3], [4], [5], [6]. Therefore, advanced understanding of the role of TAMs in cancer progression would facilitate cancer prevention, better control of disease progression, and promote the discovery of novel therapeutic approaches [7].

TAMs play a critical role in promoting tumor growth and progression through interactions with cancer cells and the TME. TAMs enhance cancer cell invasiveness by inducing epithelial-to-mesenchymal transition (EMT), a process characterized by the loss of cell adhesion and acquisition of migratory and invasive properties [8]. This is mediated by TAM-secreted factors, such as tumor necrosis factor-alpha (TNF-α) and transforming growth factor-beta (TGF-β) [8], [9]. Beyond their direct effects on cancer cells, TAMs significantly contribute to shaping a tumor-promoting microenvironment. TAMs support angiogenesis through the secretion of pro-angiogenic factors, such as vascular endothelial growth factor-A (VEGF-A) [10], [11]. This facilitates the delivery of oxygen and nutrients to the tumor mass. Within the vascularized TME, TAMs promote an immunosuppressive environment by secreting factors such as interleukin-6 (IL-6), interleukin-10 (IL-10) and transforming growth factor-β (TGF-β) in breast cancer [12], [13], [14]. TAMs also facilitate cancer-associated fibroblasts (CAFs) to promote cancer cell invasion and intravasation. This is achieved through their active participation in ECM remodeling, a critical process that enhances tumor cell dissemination and metastatic potential [15], [16].

ECM remodeling occurs through four key mechanisms: 1) ECM deposition, altering the abundance and composition of its components to modulate biochemical and mechanical properties; 2) post-translational modifications, changing the structural and biochemical characteristics of the ECM; 3) proteolytic degradation, releasing bioactive fragments and bound factors, enabling processes like cell migration; and 4) force-mediated remodeling, realigning ECM fibers and forming pathways for cellular movement. These mechanisms collectively maintain ECM dynamics, influencing tissue development, repair, and pathological conditions[17]. Although extensive research has explored TAM-mediated ECM degradation, such as MMP-9 induced collagen degradation[17], [18], [19], other aspects of ECM remodeling in response to TAMs – such as matrix synthesis, fiber crosslinking, and fiber realignment – remain poorly understood. Additionally, the murine models used in previous studies make it difficult to isolate the direct effects of TAMs on ECM structure and cancer cell behavior without the influence of fibroblasts or other stromal cells. Thus, we still lack a detailed understanding of the direct TAM-ECM interactions and their distinct roles in enabling cancer cell invasive capacity.

In this study, we employed a reductionist approach in the form of a simplified *in vitro* 3D model to study TAM-ECM interactions and their influence on cancer cell invasiveness. Our model comprised a collagen-based 3D matrix combining tumor spheroids with PBMC-derived macrophages to 1) visualize cancer cell invasion and morphological changes influenced by tumor spheroid conditioned macrophages (TSCMs) within a 3D structure, 2) characterize TSCM phenotype dynamics in the tumor microenvironment, and 3) investigate cell-ECM interactions through paracrine signaling. Using this model, we showed that breast cancer cells induced PBMC-derived macrophages to adopt a pro-tumoral phenotype. These TSCMs subsequently influenced cancer cell invasive behavior by promoting mesenchymal phenotype acquisition and mediating ECM remodeling through both collagen degradation and collagen fiber realignment.

## 2. Materials and Methods

### 2.1 Carcinoma Cell culture

MDA-MB-231 GFP breast cancer cells were obtained from the Provenzano Lab (University of Minnesota, Twin Cities) and maintained in Dulbecco’s Modified Eagle Medium (DMEM) (Gibco, Cat. # 11995-065) supplemented with 10% fetal bovine serum (FBS) (R&D Systems, Cat. # S11550) and 1% Penicillin-Streptomycin-Glutamine (Pen/strep) (Quality Biological, Cat. # 120095721) up to passage 30. BT-549 breast cancer cells were obtained from ATCC (Virginia, USA) and cultured in RPMI-1640 (ATCC, Cat. # 30-2001) supplemented with 10% FBS, 1% Pen/strep and 1 ug/mL insulin (Gibco, Cat. # 12-585-014) up to passage 30. GFP-expressing BT-549 cells were generated via stable infection with lentivirus and selected by puromycin (Gibco, Cat. # A1113803). Cells were incubated at 37°C in a humidified incubator with 5% of CO2 and fed every two days up to 80% confluence.

### 2.2 Monocyte isolation and macrophage differentiation

Buffy coats collected from healthy donors were obtained from Innovative Blood Resources (IBR) (Minneapolis, Minnesota, USA). The institutional review board (IRB) determined that the proposed activity is not research involving human subjects as defined by DHHS and FDA regulations. To arrive at this determination, the IRB used “WORKSHEET: Human Research (HRP-310).” Peripheral blood mononuclear cells (PBMCs) were isolated by density gradient centrifugation using SepMate-50 (StemCell Technologies, Cat. # 85460). Monocytes were purified from PBMCs using a monocyte enrichment negative magnetic selection bead kit (StemCell Technologies, Cat. # 19669) according to manufacturer’s instructions. Isolation with this method yielded monocytes purity over 90%, verified by flow cytometry assay.

Monocytes were cultured in ultra-low attachment 6-well plate at a density of 5x10^5^ cells/mL in RPMI media supplemented with 10% FBS, 1%Pen/strep and 60 ng/mL human macrophage colony-stimulating factor (M-CSF) (Biolegend, Cat. # 574806) for 7 days. On Day 5, without removing the culture media, same volume of fresh media supplemented with 120 ng/mL M-CSF was substituted into each well. On Day 7, to harvest cells, attached macrophages were washed by Dulbecco’s phosphate-buffered saline (DPBS) (Corning, Cat. # 21-031-CV) and incubated at 37°C in 5 mM Ethylenediaminetetraacetic acid (EDTA) (Corning, Cat. # 46-034-CI) for 30 min. After detachment, cells were washed with DPBS and centrifuged for 10 min at 400xg for following assays.

### 2.3 Tumor spheroid generation

Carcinoma cells were harvested around 70% confluence and resuspended at a density of 1x10^5^ cells/mL in culture media with 2.5% Matrigel (Corning, Cat. # 356231). 100 µL cell solution was plated into each well of an ultra-low attachment U-bottom 96-well plate (Corning, Cat. # 7007), followed by 5 min of centrifuging at 1000 rpm. Tumor spheroids were cultured at 37°C and harvested on Day 5 for following experiments.

### 2.4 Tumor invasion assay

Tumor invasion under the effect of TSCMs is evaluated in collagen I hydrogel. Rat tail collagen I (Corning, 354249) was buffered with 10x DPBS (Fisher Scientific, Cat. # BP399500), neutralized to pH 7.4 and diluted to a concentration of 2 mg/mL. Tumor spheroids harvested on Day 5 were washed by DPBS and transferred into collagen solution on ice. PBMC-derived macrophages were incubated with CellTrace FarRed (Thermo Fisher, Cat. # C34564) for 20 min at room temperature and washed by DPBS for 3 times. 1x10^5^ stained PBMC-derived macrophages were suspended in 130 uL collagen solution and kept on ice for a single sample. To ensure tumor spheroids can be fully encapsulated in 3D, 50 uL of collagen solution was added into a 96-well tissue culture plate and incubated for 5 min in the incubator, followed by 1 min incubation at room temperature. Then, an additional 80 μL of collagen solution was gently added on top of the bottom layer in a single well followed by a quick injection of a single tumor spheroid in the center of the well. After embedding, the plate was placed at 37°C for 20 minutes and fed with 180 uL 50% carcinoma cell culture media and 50% macrophage culture media supplemented with 60 ng/mL M-CSF. This embedding process was repeated for remaining wells. Each well was fed every day with respective growth media and 60 ng/ml M-CSF until Day 5. Cancer cell invasion was captured every 24 hours at 5X using a Zeiss Axio microscope with a step size of 10 μm through z-position.

### 2.5 Microscopy and imaging processing

To monitor cancer cell invasion from Day 0 to Day 5, Zeiss Axio Observer widefield microscope and a Nikon AX R confocal microscope were used. PBMC-derived macrophages were stained with CellTrace FarRed (Thermo Fisher, C34564) following manufacturer’s instructions. Collagen fibers were visualized using laser scanning multi-photon microscopy (LSMPM) and second harmonic generation (SHG). Microscope was equipped with 4-channel detection, interchangeable 440/20, 460/50, 525/50, 595/50, 605/70, and 690/50 filters (Prairie Technologies/Bruker) and a Mai Tai Ti:Sapphire laser (Spectra-Physics). PBMC-derived macrophages were stained with CellTracker Red Fluorescent Probes (Thermo Fisher, C34552) following manufacturer’s instructions. Image processing was performed in Zen software (Zeiss) and FIJI/ImageJ software. Invasion distance analysis and intensity quantification were performed using MATLAB (R2024b). Second harmonic generation (SHG) images were analyzed using the OrientationJ plugin in FIJI/ImageJ software and CT-FIRE software to assess local fiber orientation[20]. The polar histogram were generated using MATLAB (R2024b).

### 2.6 ECM dissociation and Cell sorting

Cancer cells invaded from the tumor spheroid into the collagen hydrogel as indicated above. On Day 5, TSCMs, invasive cancer cells and tumor core were extracted from the gel using 2 mg/mL collagenase (Sigma, C9263-100MG) solution. Tumor core was separated from the cell solution using 35 um cell strainers (StemCell, Cat. # 38030) and washed by DPBS. Cell solution was collected into the Falcon round bottom tube (StemCell, Cat. # 38030) and performed cell sorting at the University Flow Cytometry Resource (UFCR) of the University of Minnesota to separate the invasive cancer cells and TSCMs.

### 2.7 RNA extraction and quantitative RT-PCR assay

To characterize the phenotypes of TSCMs after 5 days of coculture with tumor spheroid in collagen ECM, RNA was isolated from cell pellets directly after cell sorting using the Single Cell RNA Purification Kit (Norgen Biotek) according to the manufacturer’s protocol. RNA concentration and quality were measured at the University of Minnesota Genomics Center (UMGC). cDNA was reverse transcribed (RT) using qScript complementary DNA (cDNA) Supermix (Quantabio, Beverly Hills, MA, USA) and then diluted with RNase-free water for mRNA amplification. qRT-PCR was performed using PowerTrack SYBR Green Master Mix (Thermo Fisher Scientific, Cat. # A46113) and the QuantStudio 3 Real-Time PCR system. Gene expression was calculated using the efficiency corrected method (ΔΔCt) and normalized to the housekeeping gene Glyceraldehyde-3-phosphate dehydrogenase (GAPDH). All primers (GAPDH, CD163, CD206) were purchased from Sigma-Aldrich. Primer sequences used for qRT-PCR: GAPDH F: GTCTCCTCTGACTTCAACAGCG; GAPDH R: ACCACCCTGTTGCTGTAGCCAA; CD163 F: CCAGAAGGAACTTGTAGCCACAG; CD163 R: CAGGCACCAAGCGTTTTGAGCT; CD206 F: AGCCAACACCAGCTCCTCAAGA; CD206 R: CAAAACGCTCGCGCATTGTCCA.

### 2.9 Immunofluorescent (IF) staining

TSCMs were harvested from the gel as described above and plated into 96-well tissue culture plate. 96-well plate was incubated at 37°C for two hours to allow cell attachment. 4% paraformaldehyde (PFA) solution was used to fix cells. 5% BSA (Sigma, A9418) and 0.1% Tween 20 (Sigma, P5927) in DPBS solution was used to block and permeabilize macrophages. Mouse anti-human CD163 primary antibody (Thermo, Cat. # TA506391), rabbit anti-human CD206 primary antibody (Thermo, Cat. # MA5-32498), goat anti-mouse 555 secondary antibody (Thermo, Cat. # A-21422), donkey anti-rabbit 647 secondary antibody (Thermo, Cat. # A-31572), Hoechst (Thermo, Cat. # H3569) were diluted in 1% BSA/DPBS at 1:100, 1:100, 1:200, 1:200 and 1:1000, respectively. Macrophages were incubated with primary antibodies for 1 hour at 37°C, followed by DPBS washing for 3 times. Macrophages were then incubated with secondary antibodies and Hoechst for 1 hour at 37°C, followed by DPBS washing for 3 times. Fluorescent images were captured at 10X using a Zeiss Axio microscope.

### 2.10 Proteome profiler assay

To collect the soluble factors secreted by tumor spheroid and macrophages from the collagen hydrogel, the supernatant was collected and spun at 400xg for 10 mins at 4 °C to remove cell debris. This supernatant was used to a Proteome Profiler Human Oncology Array (R&D Systems, Cat. # ARY026). All steps of the secretion assay were performed according to the manufacturer’s protocol. Protein expression was normalized by the total protein amount in the conditioned media. The total protein amount was measured using BCA protein assay. Expression of soluble factors was analyzed by densitometry using ImageStudio Lite 2.2 (Licor).

## 3 Statistical Methods

All quantitative values represent mean ± standard error of at least three independent experiments. Comparison between experimental groups of equal variances were analyzed using an unpaired t-test and 95% confidence interval or one-way ANOVA followed by Tukey’s test for multiple comparisons. Statistical calculations were performed using GraphPad Prism 10.0. The difference was considered significant at P < 0.05 (*p < 0.05, **p < 0.01, ***p < 0.001, ****p < 0.0001).

## 4 Results

### 4.1 3D model enables quantification of cancer cell invasiveness into the TME

Tumor spheroid models provide a 3D structure to study cell-cell and cell-ECM interactions. MDA-MB-231 is a highly aggressive human triple negative breast cancer cell line in oncology studies [21]. It shows representative EMT progress associated with cancer progression. In our study, we used MDA-MB-231 as the tumor model to study breast cancer invasion. Specifically, the large-scale tumor spheroid was formed by seeding 10,000 MDA-MB-231 GFP breast cancer cells in a round bottom ultra-low adhesion well plate. Representative images of C tumor spheroids during the first 5 days demonstrated the spontaneous formation of compact tumor spheroids and the increase of tumor size over time (Fig S1a-f). On Day 5, we harvested the MDA-MB-231 GFP tumor spheroids with the diameter over 500 um (Fig. 1a) and embedded tumor spheroids into collagen I hydrogel, as collagen I is a crucial component of the ECM in the TME and strongly associated with tumor growth, invasion and metastasis (Fig. 1b). Cancer cells invaded from the tumor spheroid were visualized in this optically accessible model.

**Figure 1.**
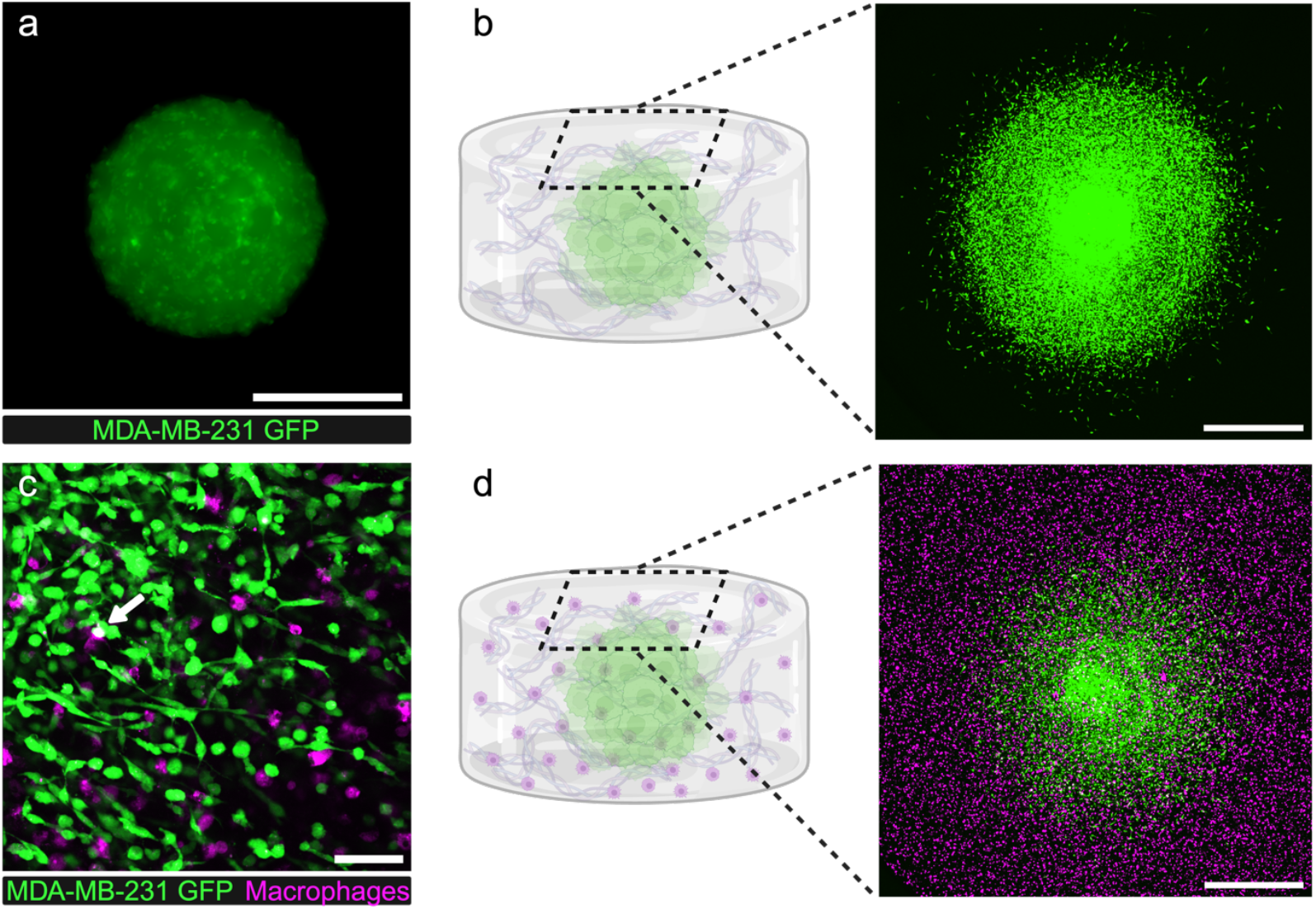
Collagen-based tumor spheroid model with PBMC-derived macrophages. (a) Fluorescence image of an MDA-MB-231 GFP tumor spheroid seeded with 10,000 cancer cells and cultured for 5 days (Scale bar: 500 μm). (b, d) Schematic representation of collagen-based tumor spheroid model without macrophages (b) and with macrophages (d) and representative confocal image after orthogonal projection showed invasive cancer cells (green) and surrounding macrophages (magenta) for each setup on Day 5 (Scale bar, 500 μm). (c) Representative multiphoton image of tumor margin with invasive cancer cells (green) and macrophages (magenta) on Day 5 (Scale bar, 100 μm). Colocalization was labeled by the white arrow.

It has been previously documented that TAMs support tumor progression by promoting cancer proliferation, invasion and metastasis[22], [23], [24]. In our study, to investigate the effects of TAMs on cancer invasion within the collagen matrix, unstimulated PBMC-derived macrophages were introduced into the 3D tumor environment. Specifically, 100,000 PBMC-derived macrophages were embedded in the collagen matrix of each well, with an MDA-MB 231 tumor spheroid positioned at the center and co-cultured for five days (Fig. 1d). During the five-day culture period, MDA-MB 231 cancer cells progressively invaded into the surrounding collagen matrix, leading to a reduction in the distance between cancer cells and macrophages. By later time points, colocalization of cancer cells and macrophages was observed (Fig. 1c). This optically accessible model enabled visualization of cancer cells and macrophages in the 3D structure, which provided a platform to quantify cancer cell invasion and morphology in response to macrophages in the 3D TME.

### 4.2 PBMC-derived macrophages promote a shift in cancer cell populations toward a more invasive phenotype, characterized by increased invasion distance

We hypothesized that PBMC-derived macrophages in the TME can promote cancer cell invasion. Previous studies have shown that PBMC-derived monocytes can be recruited into the TME and differentiate into TAMs [25], [26]. TAMs play a critical role in promoting tumor growth and progression through interactions with cancer cells and the TME. Figure 2a showed MDA-MB 231 tumor spheroid invasion in the 3D collagen hydrogel and visualized the influence of PBMC-derived macrophages on cancer invasion. Specifically, in the tumor spheroid without macrophages (TS^+^/Mφ^-^) condition,

**Figure 2.**
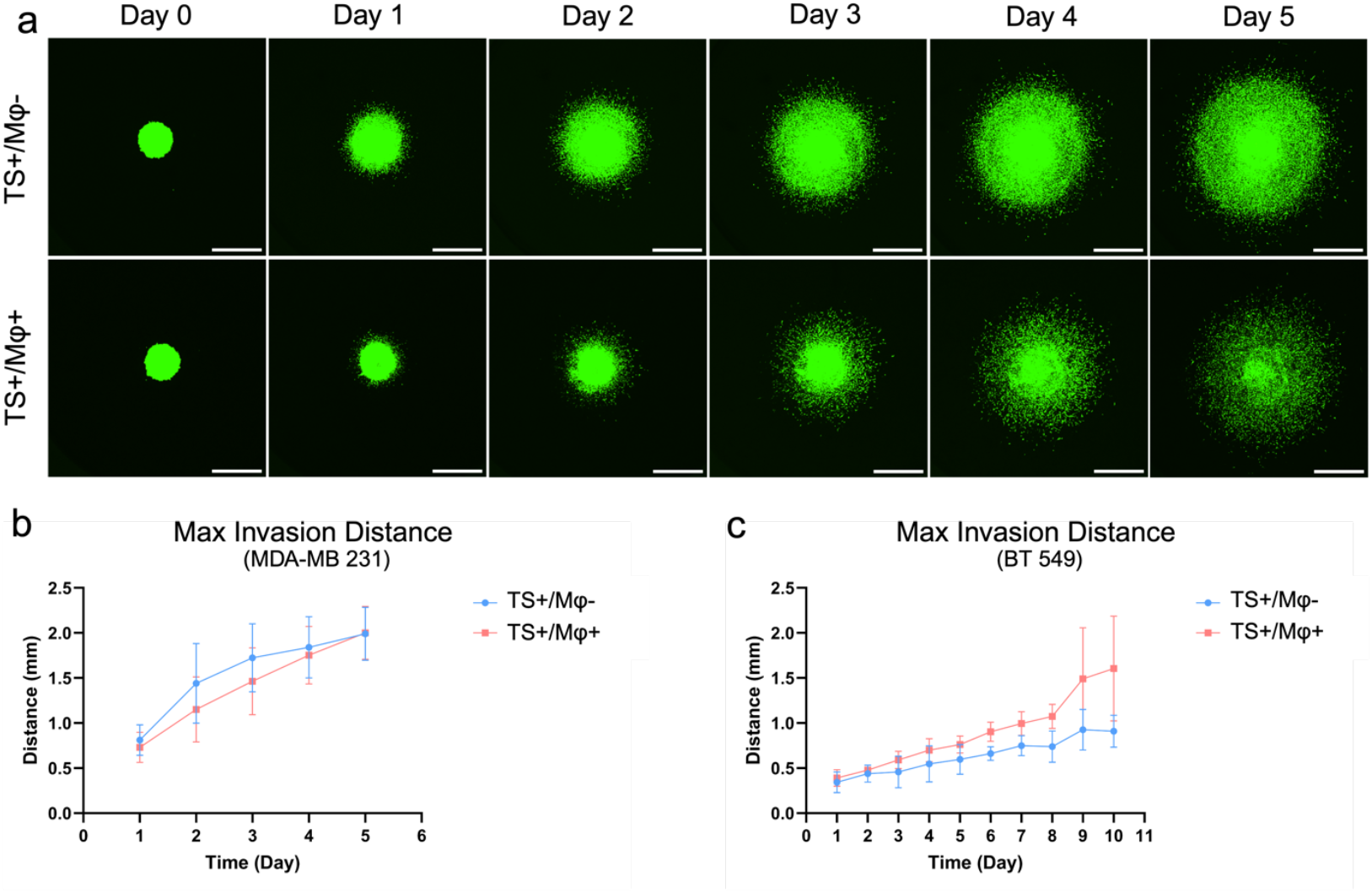
Visualizing and quantifying of cancer invasion responding macrophages. (a) Representative confocal images of tumor invasion from Day 0 to Day 5 in the TS^+^/ Mφ^+^ condition and the TS^+^/ Mφ^-^ condition (Scale bar: 500 μm). (b) MDA-MB-231 GFP cancer cell maximum invasion distance every 24 hours in the TS^+^/ Mφ^+^ condition and the TS^+^/ Mφ^-^ condition during the first five days. (c) BT-549 GFP cancer cell maximum invasion distance every 24 hours in the TS^+^/ Mφ^+^ condition and the TS^+^/ Mφ^-^ condition during the first ten days.

MDA-MB-231 GFP cancer cells invaded outward from the tumor spheroid into the surrounding collagen ECM, forming a high intensity ring at the periphery, indicating high cell density. Beyond this high-density region, a small number of cancer cells exhibited further invasion. By contrast, no high-density ring region was not observed in the tumor spheroid with macrophages (TS^+^/ Mφ^+^). Instead, invasive cancer cells were more evenly distributed along the radius in the surrounding collagen ECM. Interestingly, cancer cells in the TS^+^/Mφ^-^ condition exhibited higher proliferation indicated by higher GFP intensity compared to the TS^+^/ Mφ^+^ condition. Although TAMs can support tumor progression, including cancer cell proliferation and invasion, macrophages in our model on the early stage appear to inhibit cancer cell proliferation. This could be explained by the absence of tumor pre-conditioning of the macrophages in our model, which could lead to the release of inflammatory cytokines that impede cellular proliferation in the first few days.

To investigate the impacts of PBMC-derived macrophages on MDA-MB 231 cancer cell invasion, we quantified the maximum invasion distance every 24 hours during the first five days. The TS^+^/Mφ^-^ condition showed an increased maximum invasion distance characterized by a rapid initial increase prior to Day 3, followed by a slower expansion of the invasive front beyond Day 3. By contrast, the TS^+^/ Mφ^+^ condition exhibited a more consistent, linear trend with a steady slope. On Day 5, the maximum invasion distance of the TS^+^/ Mφ^+^ condition reached the TS^+^/Mφ^-^ condition (Fig 2b). Considering the steeper slope of the curve observed in the later stage of TS^+^/ Mφ^+^ condition, it was expected that the maximum invasion distance in this condition would be higher than that of the TS^+^/Mφ^-^ condition after Day 5. However, due to the high invasiveness of MDA-MB-231 GFP cancer cells, the most distant cells had already reached the boundary of the tissue culture plate, making it impossible to track cancer invasion beyond Day 5. Although the MDA-MB-231 line is one of the most common triple-negative breast cancer cell lines used in oncology studies, its highly proliferative and malignant characteristics may obscure the pro-tumoral effects from macrophages.

To test the effect of PBMC-derived macrophages on the invasiveness of a less proliferative and invasive cell line, we used BT-549, another human triple negative breast cancer cell line. Figure 2c showed that the invasion distance of BT-549 GFP cancer cells in the TS^+^/ Mφ^+^ condition is higher than that in the TS^+^/ Mφ^-^ condition, especially on later days when macrophages had been exposed to the tumor spheroids for an extended period. Figure S2 illustrated that a greater number of cancer cells from BT-549 GFP tumors invaded into the collagen ECM on Day 10 in response to macrophages. These findings demonstrated that PBMC-derived macrophages in the tumor environment can modulate cancer cell invasion behaviors.

### 4.3 PBMC-derived macrophages modulate cancer cell morphology in collagen matrix

Cell shape is closely associated with cancer cell malignancy [27], [28]. Cancer cells become elongated with reduced cell-cell adhesion during the EMT [29], [30]. The longer protrusions can facilitate cell movement along the ECM network and consequently promote cancer cell invasion in the surrounding ECM. Thus, to further understand the impact of PBMC-derived macrophages on cancer cell invasion, we evaluated MDA-MB-231 GFP cancer cell morphology at the tumor margin, where cancer cells were exposed to high concentrations of macrophage-secreted factors and may engage in cell-cell interactions within the collagen hydrogel containing macrophages. Figure 3a-b showed that MDA-MB-231 GFP cancer cells in the TS^+^/ Mφ^-^ condition exhibited more circular morphologies, whereas cells in the TS^+^/ Mφ^+^ condition displayed a shift toward an elongated phenotype, as quantified by cell circularity (Fig. 3c). This lower circularity of cancer cells in the TS^+^/ Mφ^+^ condition may reflect the cytoskeleton reorganization of cancer cells in response to macrophages. We also measured the impact of PBMC-derived macrophages on cancer cell morphology in BT-549 GFP cancer cells. Consistent with the findings in MDA-MB-231 GFP cancer cells, Figure S2 demonstrated that a higher proportion of BT-549 GFP cancer cells displayed a spindle-like morphology in the TS^+^/Mφ^+^ condition and the cell circularity of BT-549 GFP cancer cells in the TS^+^/ Mφ^+^ condition is significantly lower than that in the TS^+^/ Mφ^-^ condition (Fig 3d).

**Figure 3.**
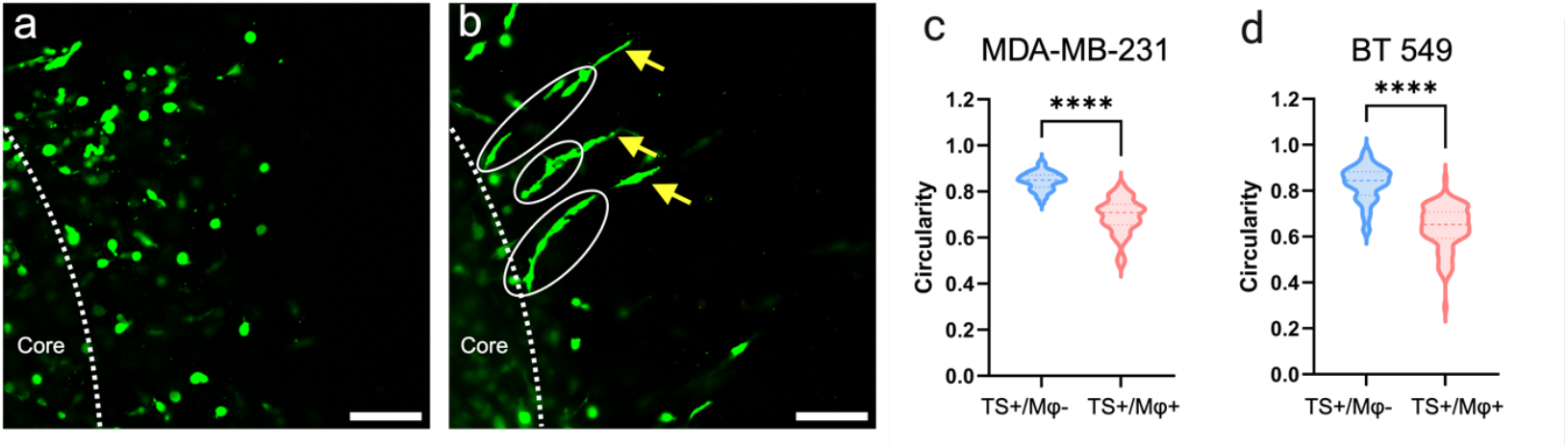
Characterization of cancer cell morphology in collagen hydrogel. (a) Representative image of MDA-MB-231 GFP tumor margin in the TS^+^/ Mφ^-^ condition with higher circularity invasive cancer cell (Scale bar: 100 μm). (b) Representative image of MDA-MB-231 GFP tumor margin in the TS^+^/ Mφ^+^ condition with collective invasion cancer cells. Leader cancer cells were pointed by yellow arrows and the follower clusters were labeled by white ovals (Scale bar: 100 μm). (c-d) Quantification of MDA-MB-231 GFP (c) and BT-549 GFP (d) cell circularity in ROIs. Ten ROIs were randomly picked from one image and the cell circularity from the ten ROIs

Moreover, collective cancer cell invasion with leader-follower organization was frequently observed in TS^+^/ Mφ^+^ condition (Fig. 3a-b, S2a-b). Leader cells directed the invasion of the clusters, with follower cells invading sequentially behind the leader cells. In the clusters, some follower cells were connected to neighboring cells via protrusions, while others were disconnected but remained aligned within the invasion track directed by the leader cells (Fig. 3b, S2b). This disconnection may suggest that the leader-follower organization operated in a cell contact-independent manner. Cells within the invasion clusters exhibited low circularity and extended long protrusions into the surrounding ECM (Fig. 3b, S2b). These findings provided further evidence that PBMC-derived macrophages influence cancer cell morphology at the tumor margin and highlight their role in enhancing tumor invasion efficiency.

### 4.4 Tumor spheroids promote polarization of macrophages towards a pro-tumoral M2-like phenotype

Macrophages are heterogeneous with phenotypes that have been previously described as the M1-M2 paradigm, where M1 macrophages are pro-inflammatory while M2 macrophages are reported to promote cell proliferation and tissue repair [31]. cMacrophages in the tumor microenvironment were thought to be polarized to an M2-like phenotype, due to their ability to support tumor growth, immune evasion, and metastasis [6], [32]. To investigate the crosstalk between cancer cells and macrophages and to elucidate the pro-tumoral effects on cancer invasion and cell morphology, we characterized the phenotype of macrophages in the spheroid model. In the TS^-^/ Mφ^+^ condition, unstimulated PBMC-derived macrophages were encapsulated in collagen hydrogel. While in the TS^+^/ Mφ^+^ condition, an equal number of PBMC-derived macrophages were incorporated in collagen hydrogel on Day 0 and cocultured with the tumor spheroid for 5 days under identical conditions. On Day 5, we isolated the TSCMs, which were conditioned by the tumor spheroid for five days, from the collagen hydrogel to characterize their phenotype (Fig. 4a). Isolated TSCMs were re-cultured in a 2D environment and subjected to immunofluorescence (IF) staining for CD163 and CD206, which are well-established markers of M2-like macrophages (Fig 4b). The results revealed an increased expression of CD163 and CD206 in the TS^+^/ Mφ^+^ condition, suggesting tumor spheroid conditioning shifting PBMC-derived macrophages toward pro-tumoral phenotypes. To further quantify the phenotypic dynamics of macrophages, the expression of CD163 and CD206 in the TS^+^/ Mφ^+^ condition was compared to the TS^-^/ Mφ^+^ condition using qRT-PCR (Fig 4c-d). Elevated expression of CD206 in the TS^+^/ Mφ^+^ condition was consistent with the IF staining. The expression of CD163 was not statistically different from the TS^-^/ Mφ^+^ condition, but the upward trend was observed. These results indicated that tumor spheroids stimulated PBMC-derived macrophages to the M2-like phenotype in our model, which may exhibit pro-tumoral effects on the tumor spheroids.

**Figure 4.**
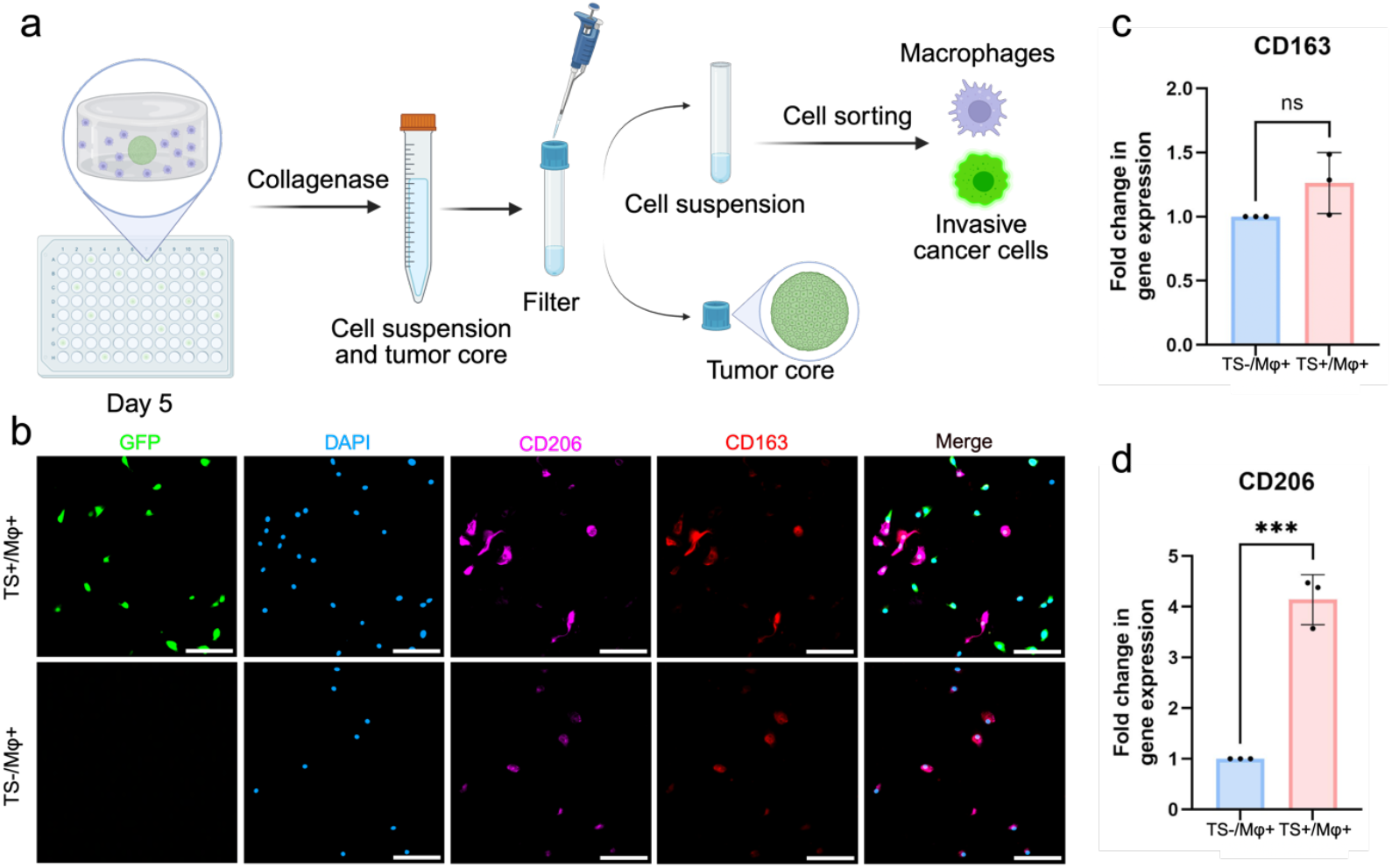
Characterization of macrophage phenotypes under the stimulation of MDA-MB-231 GFP cancer cells on Day 5. (a) Schematic representation of the pipeline to separate cellular components in the TS^+^/Mφ^+^ condition. (b) Wide field images of MDA-MB-231 GFP cancer cells Green) and macrophages stained with DAPI (nuclei, blue), CD206 (magenta) and CD163 (red) (Scale bar: 100 μm). (c-d) Gene expressions of CD163 (c) and CD206 (d) on macrophages in the TS^+^/Mφ^+^ condition and TS^-^/Mφ^+^ condition on Day 5. (n=3)

### 4.5 TSCMs promote cancer invasion through ECM remodeling

TAMs, as key stromal cells, are known to mediate ECM remodeling. To investigate the impacts of the MDA-MB-231 GFP TSCMs on the surrounding ECM, collagen degradation assays were conducted to evaluate the degradation properties of the collagen matrix. Specifically, collagen gels with tumor spheroid and macrophages were transferred to 500 uL 2 mg/mL collagenase solution and incubated at 37°C without any mechanical force. Collagen gel integrity was observed and recorded every 5 min until the gels were fully digested. Figure 5a showed that the collagen gels in the TS^+^/ Mφ^+^ condition was fully digested within 15 minutes, whereas the gels in the TS^+^/ Mφ^-^ condition remained intact and the complete digestion required an average of 30 to 35 minutes (Fig. S3). This observation indicated that TSCMs promoted collagen degradation, leading to the formation of a more fragmented ECM network. Previous studies have demonstrated that ECM degradation can release bioactive fragments and bound factors, which play a role in promoting cell migration within the TME. While our *in vitro* platform is not capable of fully elucidating this mechanism, our findings highlight the impact of TSCMs on ECM degradation.

**Figure 5.**
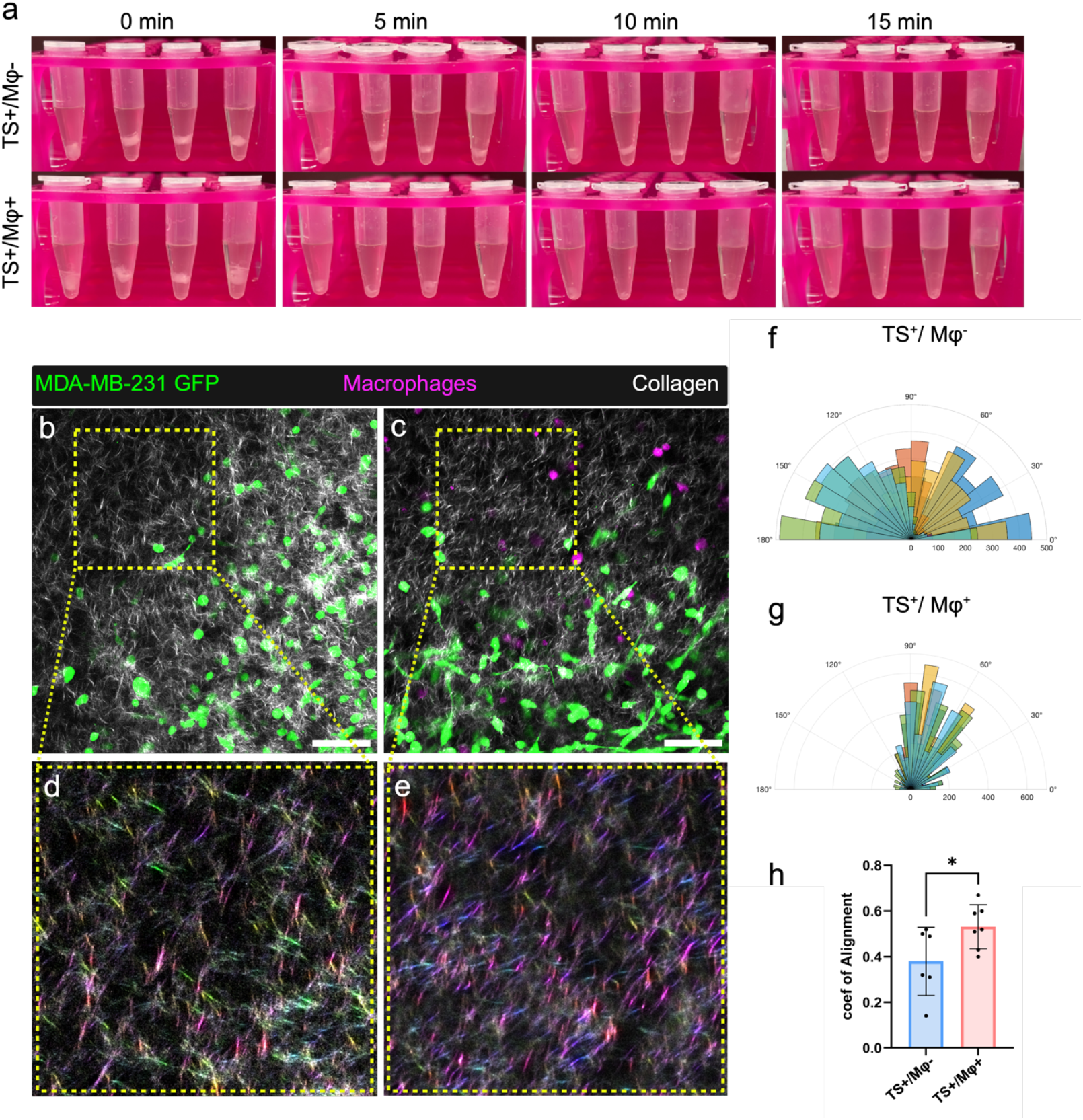
Characterization of collagen hydrogel. (a) Collagen hydrogel digestion process in the collagenase solution. The four tubes in each photo were four replicates. (b-c) Representative multiphoton images of collagen fiber (white) architecture in the TS^+^/ Mφ^+^ condition (b) and the TS^+^/ Mφ^-^ condition (c) (Scale bar: 100 μm). (d-e) Corresponding ROIs of the collagen fiber orientation from the TS^+^/ Mφ^+^ condition (b) and the TS^+^/ Mφ^-^ condition (c) using the OrientationJ plugin (ImageJ, NIH). The color map represents the angle with the horizontal axis. (f-g) Quantification of collagen fiber orientation using normalized polar histogram derived from second harmonic generation (SHG) images. Each color represents one sample (n>3). To normalize the data, all angles from each sample were subtracted by the corresponding mean value. (h) Coefficient of collagen fiber distribution.

According to previous studies, ECM remodeling resulted in distinct patterns of collagen orientation and alignments, facilitating local cancer cell invasion into surrounding ECM, termed Tumor-Associated Collagen Signature (TACS) [33], [34]. In our study, we investigated the remodeling of collagen network mediated by TSCMs in our cancer invasion model, collagen fibers were visualized using laser scanning multiphoton microscope. In the TS^+^/ Mφ^-^ condition, collagen fibers appeared disorganized and lacked directional alignment, indicative of a less structured extracellular matrix architecture in the absence of TSCMs (Fig. 5b). However, in the TS+/Mφ+ condition we observed prominent collagen fiber alignment extending radially from the tumor boundary, suggesting TSCM-driven matrix organization (Fig. 5c). To further visualize the differences in collagen fiber orientation, representative OrientationJ-generated orientation colormaps were analyzed. As shown in Figure 5d–e, the TS+/Mφ − condition displayed a broader distribution of colors in the colormap, whereas the TS+/Mφ+ condition exhibited relatively uniform fiber alignment with minimal color variation, suggesting a more organized collagen network.

The collagen fiber distribution was quantified using CT-FIRE analysis [20]. The polar histograms (Fig. 5g-f) show a narrower distribution of collagen fiber angles in the TS^+^/ Mφ^+^ condition than in the TS^+^/ Mφ^-^ condition, which indicating a more organized collagen network in the presence of macrophages. The coefficient of alignment of the TS^+^/ Mφ^+^ condition was significantly higher than the TS^+^/ Mφ^-^ condition (Fig. 5h). The higher kurtosis, lower standard deviation and variance further demonstrated the greater uniformity of collagen fiber angle distribution in the TS^+^/ Mφ^+^ condition (Fig. S4a-c). The aligned collagen fibers can create tracks to guide the migration of cancer cells, promoting invasion and metastasis. Combining the previous findings on collagen degradation, our study suggested that TSCMs promoted cancer cell invasion through ECM remodeling. TSCM-induced ECM degradation generated weaker ECM and therefore may reduce resistance to cancer cell movement through the 3D matrix. The aligned collagen fibers provided invasion tracks, enabling more efficient navigation of cancer cells through the ECM and enhancing invasive capacity.

### 4.6 Soluble factor profile analysis in the tumor invasion model with TSCMs

To understand the cellular and molecular mechanisms of TSCM-induced ECM remodeling, secretome analysis was performed using conditioned media from the TS^+^/ Mφ^+^ condition, TS^+^/ Mφ^-^ condition and TS^-^/ Mφ^+^ condition. MDA-MB-231 GFP cancer cells were used to generate the tumor spheroid. The conditioned media were collected on Day 5 and the analysis was performed using human oncology proteome profiler array (Fig. S5). Our analysis revealed that MMP-9 is highly enriched in the TS^+^/ Mφ^+^ condition and the TS^-^/ Mφ^+^ condition, which is consistent with our observation that collagen gels from the TS^+^/ Mφ^+^ condition degraded faster than the TS^+^/ Mφ^-^ condition. Surprisingly, the levels of MMP-9 in the TS^-^/ Mφ^+^ condition were slightly higher than that in the TS^+^/ Mφ^+^ condition, which may result from the suppression of cancer cells on MMP-9 production by macrophages (Fig 6a). Cathepsins B, D, and S, as important proteases to promote tumor invasion, directly degrade ECM components, such as collagen, and indirectly promote ECM degradation by activating MMPs [35], [36], [37], [38]. These factors were upregulated in the TS^+^/ Mφ^+^ condition, which may directly facilitate collagen degradation to provide a loosened network for cancer cell migration (Fig 6b-d).

**Figure 6.**
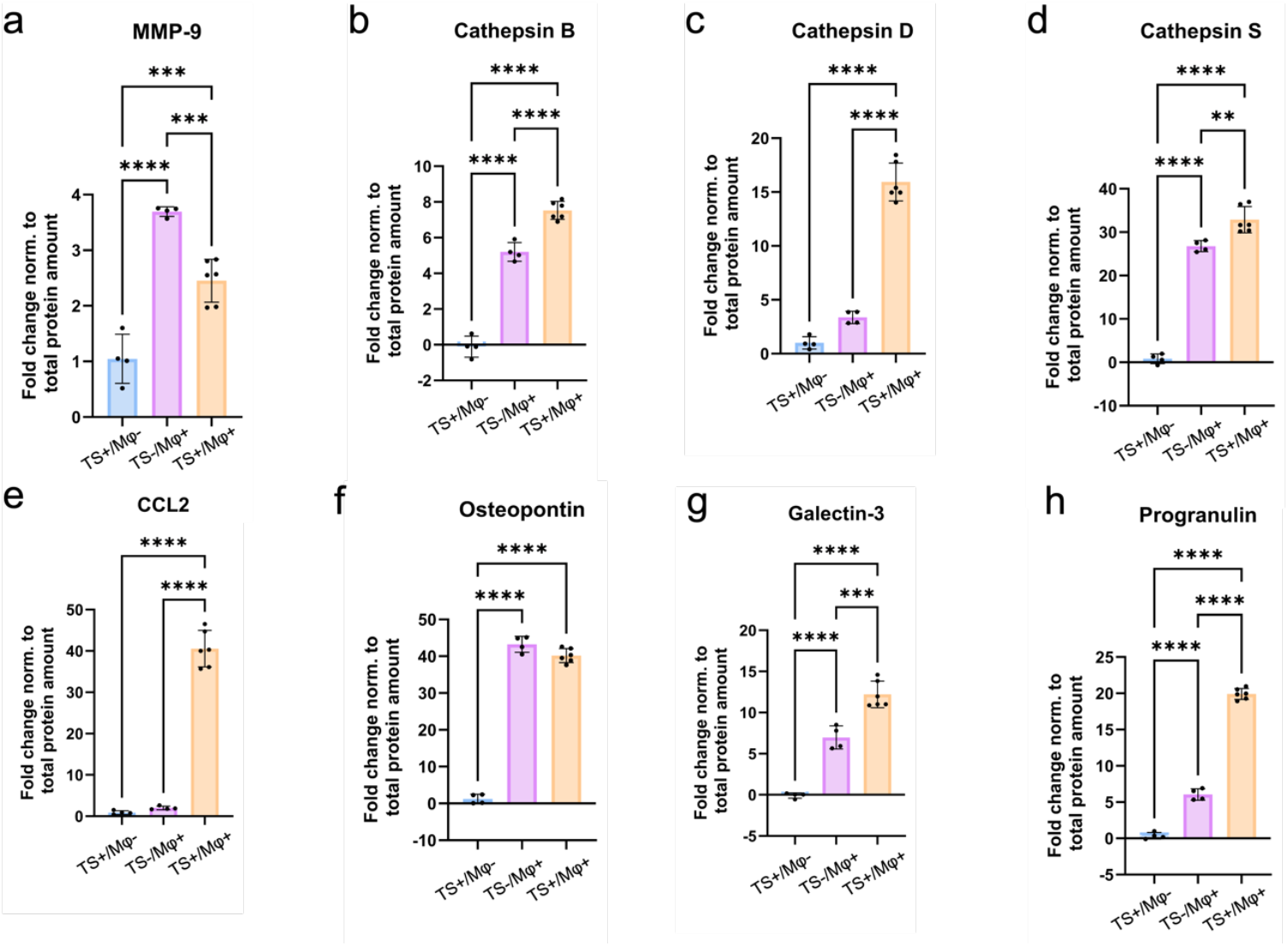
Secretome analysis of tumor related proteins in the tumor invasion model as revealed by a human oncology proteome profiler array. (a-h) Quantification of tumor related protein levels from the conditioned media in the TS^+^/ Mφ^+^ condition, TS^-^/ Mφ^+^ condition and TS^+^/ Mφ^-^ condition. (n >3)

CCL2 was significantly upregulated in the TS^+^/ Mφ^+^ condition compared to the other two conditions, which further demonstrated the PBMC-derived macrophages were polarized to the pro-tumoral phenotypes in our model (Fig. 6e). CCL2 can upregulate the expression of MMPs and stimulate macrophages to secrete more cathepsins and consequently contribute to ECM remodeling [39]. Other soluble factors that were upregulated in the TS^+^/ Mφ^+^ condition compared to TS^+^/ Mφ^-^ condition, such as Osteopontin (OPN), Galectin-3 and Progranulin (PGRN), indirectly contributed to ECM degradation by activating MMP-2 and MMP-9 (Fig. 6f-h). According to previous studies, these soluble factors can promote collagen degradation to create a low-resistant ECM and may consequently facilitate cancer cell invasion [40], [41]. Additionally, OPN, Galectin-3 and PGRN can promote EMT by downregulating the expression of epithelial markers, such as E-cadherin, and upregulating mesenchymal markers, such as N-Cadherin and vimentin [42], [43], [44]. OPN and Galectin-3 also facilitate cancer cell detachment from the tumor mass and promote cancer cell adhesion to ECM [42], [43], [45]. OPN induced cytoskeletal reorganization [45], which is consistent with the observed changes in cell circularity in our study. Therefore, the upregulation of OPN, Galectin-3 and PGRN in the condition with TSCMs can promote cancer cell invasion ability and indirectly contribute ECM remodeling to facilitate cancer migration.

## 5 Discussion

In our study, we used a collagen-based 3D tumor spheroid model to study the interactions among cancer cells, macrophages, and collagen ECM. Our findings demonstrated that cancer cells shifted macrophage population toward the M2-like macrophage phenotype. In turn, such pro-tumoral TSCMs drove a shift in the cancer cell population toward a phenotype with greater invasive capacity and lower cell circularity, which indicated higher aggressiveness. Additionally, the pro-tumoral TSCMs modulated collagen degradation and fiber realignments to provide a tumor-favoring microenvironments for cancer cell invasion. These findings emphasized the phenotypical dynamics of macrophages in the TME and the impacts of pro-tumoral macrophages on cancer progressions through ECM remodeling. Having the capacity to understand the cancer cell-macrophage-ECM interactions, this 3D tumor model has the potential to reveal predictive bio-targets and prognostic indicators for cancer therapies.

Cancer cells acquire a spindle-like or irregular morphology, reflecting increased plasticity and adaptability. Such cell morphological changes support cancer cells to invade in tissue and navigate complex microenvironment. EMT, as a critical process in cancer progression, induces the transition of high circularity to low circularity, which shift cancer cells toward a more aggressive, motile, and invasive behavior. Our study revealed that TSCMs modulated cancer cell circularity, indicating the promotion of mesenchymal phenotype acquisition. Additionally, cancer cells orient their migration in response to mechanical and biochemical properties in the TME, including neighboring cells and ECM. In collective cell migration, such phenomenon can facilitate the coordination between leader and follower cells. In our study, such leader-follower organization was frequently observed in the TS^+^/ Mφ^+^ condition. Leader cancer cells responded to the soluble factor signals and mechanical forces from surrounding collagen ECM under the impacts of TSCMs, creating cohesive invasion fronts. Follower cells detected and responded to the leader cells within the collective cell cluster, resulting in enhanced migratory efficiency. These findings offered novel insights into the role of TSCMs in regulating cancer cell invasive capacity and their contribution to tumor progression.

Previous studies revealed that TAMs are prominent producers of MMPs, including MMP-2 and MMP-9, which degrade collagen and other ECM components, and consequently supporting a microenvironment conducive to cancer progression[46]. Consistent with these findings, our results highlighted the role of macrophages in MMP secretion. Additionally, our work expanded the analysis of the secretome and identified additional factors, including cathepsin B, D, S, CCL2, OPN, galectin, and PGRN, which were upregulated in the TS^+^/ Mφ^+^ condition. These factors have been previously shown to directly or indirectly contribute to ECM degradation. Notably, the level of CCL2 was highly upregulated in the TS^+^/ Mφ^+^ condition compared to the TS^+^/ Mφ^-^ condition and the TS^-^/ Mφ^+^ condition. In addition to its contribution to ECM degradation, CCL2, as a chemokine, can recruit monocytes to the TME in *in vivo* models, where the monocytes differentiated to macrophages, which can develop a positive feedback loop to facilitate cancer invasion. The upregulation of these factors in our model may offer a cellular and molecular basis for understanding how TSCMs promote cancer invasion within the collagen hydrogel system through ECM degradation.

In addition to promoting ECM degradation, our study highlighted collagen fiber realignment in the TME with macrophages. This observation provides a novel perspective on TSCM-mediated ECM dynamics, challenging the previous notion that fiber realignment is predominantly driven by fibroblasts. However, the precise mechanism underlying macrophage-induced collagen fiber realignment remains unclear in our study. Macrophages may physically alter the organization of collagen fibers by exerting forces during their migration through the collagen matrix. Some macrophage-secreted cytokines and soluble factors may indirectly influence collagen alignment by modulating ECM components, although these effects are not as strong as the functions of fibroblasts. Another potential mechanism involves the macrophage-secreted soluble factors, such as MMPs and Cathepsins, which contribute to ECM degradation. In our study, the collagen gels were demonstrated to be weaker in the TS^+^/ Mφ^+^ condition. Therefore, during the process of cancer cells invade from the tumor into the surrounding ECM, the mechanical force of cancer cells may stretch and realign the weaker collagen gel, creating an aligned matrix structure. This aligned ECM could subsequently serve as a migration track for cancer cells, enhancing their invasive capability. Therefore, the mechanisms of the collagen fiber realignment require further study.

Taken together, our study used a 3D tumor spheroid model to study cancer cells behaviors in the 3D structure under the impact of TSCMs. We elucidated that TSCMs modulated cancer invasion through ECM remodeling. Additional studies are needed to identify signaling pathways involved in these soluble factors and cell-cell, cell-ECM interactions, such as the role of CCL2 in ECM remodeling in addition to immune cell recruitment, integrin-mediated cell adhesion in collagen ECM during cancer cell migration process, leader-follower organization in the 3D ECM and mechanism underlying macrophage-mediated collagen fiber realignment. In this work, we mainly used MDA-MB-231 GFP cancer cell line to investigate the interactions among tumor spheroids, macrophages, and surrounding ECM, and used BT-549 GFP cells to validate the invasion and cancer cell morphology phenotype. Future investigations can also aim to incorporate more cellular components and clinically relevant structures to better recapitulate the physiological relevant process. Ultimately, it is essential to use an optically accessible 3D *in vitro* model to recapitulate the complex TME with cellular and non-cellular components and provide robust platform to advance our understanding of cancer progression and explore personalized treatment approaches.

## 6 Conclusion

In this study, we used a 3D collagen-based tumor spheroid model to investigate the impact of PBMC-derived macrophages on breast cancer invasion. Our findings indicated that tumor spheroid stimulated PBMC-derived macrophages into a pro-tumoral phenotype, and the TSCM induced a shift in cancer cell populations toward a phenotype characterized by increased invasion distance, while also modulating cancer cell morphology within the 3D collagen hydrogel. Collagen degradation assays and fiber characterization provided evidence that TSCMs in the tumor microenvironment promoted cancer invasion through ECM degradation and collagen fiber realignment. Additionally, secretome analyses identified alterations in soluble factors that play a critical role in ECM remodeling within the system. In conclusion, our study advanced the understanding of macrophage-mediated ECM remodeling and the influence on tumor invasion using an optically accessible 3D *in vitro* model.

## CRediT authorship contribution statement

**Luna Zhang:** Conceptualization, Data curation, Formal analysis, Investigation, Methodology, Project administration, Software, Visualization, Writing – original graft. **Sanab Abdi:** Data curation, Formal analysis.

**Hannah M. Szafraniec:** Conceptualization, Formal analysis.

**Paolo P. Provenzano:** Conceptualization, Resources.

**Kathryn L. Schwertfeger:** Conceptualization, Validation, Writing – review & editing, Supervision, Resources.

**David K. Wood:** Conceptualization, Validation, Writing – review & editing, Supervision, Resources, Project administration, Funding acquisition.

## Acknowledgements

Support for this work was provided by the NCI under R01CA245550, U54CA210190, P01CA254849, R01CA215052, R01CA265004 and the Interdisciplinary Doctoral Fellowship (IDF) from the University of Minnesota. Portions of this work were conducted in the Minnesota Nano Center, which is supported by the National Science Foundation through the National Nanotechnology Coordinated Infrastructure (NNCI) under Award Number ECCS-2025124.

The authors also wish to thank the Provenzano Lab at the University of Minnesota, especially Demi (Hongrong) Zhang and Guhan Qian for their help with using multiphoton microscopy and image analysis. The Schwertfeger Lab is warmly acknowledged for their technical assistance on biological assays. The authors gratefully acknowledge Emilio Tarcitano, PhD, Elizabeth L. Crist, PhD and Maggie Chiu for their technical guidance.

## Appendix. Supplementary materials

**Figure S1.**
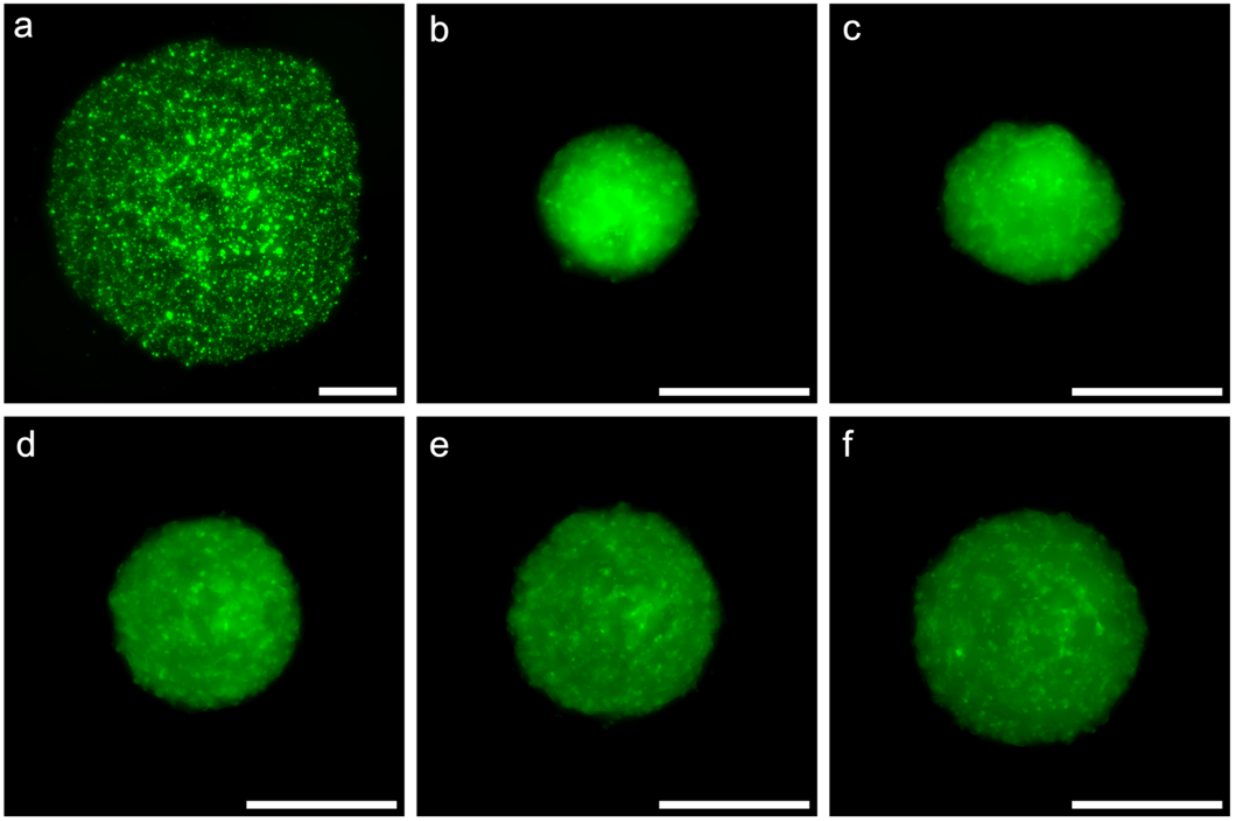
The growth of tumor spheroid. (a-f) MDA-MB-231 GFP tumor spheroid from Day 0 to Day 5 (Scale bar: 500 μm).

**Figure S2.**
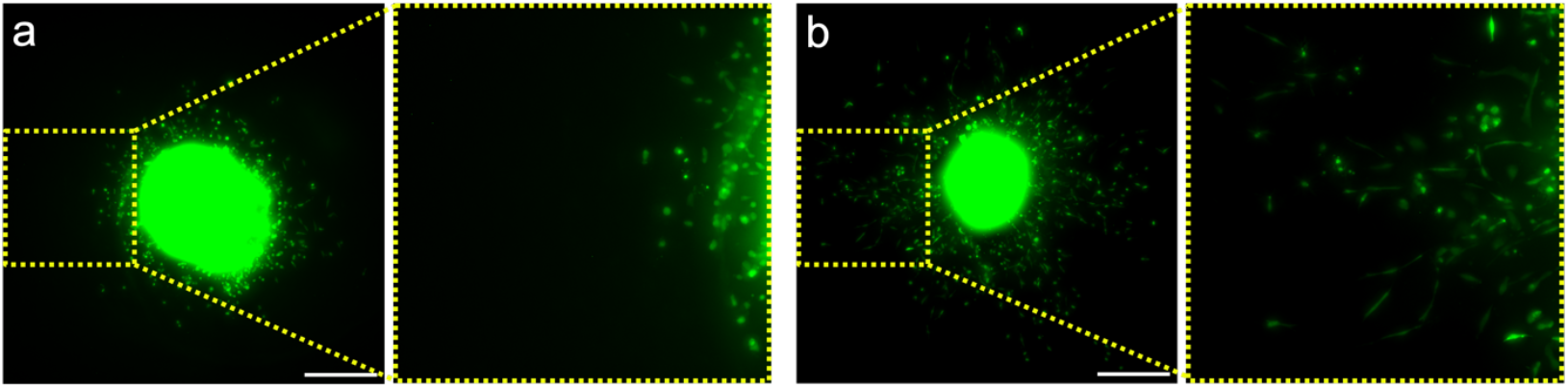
Tumor invasion and cell morphology of BT-549 GFP. (a-b) Representative wide field images of BT-549 GFP tumor spheroid invasion on Day 10 in the TS^+^/ Mφ^-^ condition (a) and the TS^+^/ Mφ^+^ condition (b). Scale Bar: 500 um.

**Figure S3.**
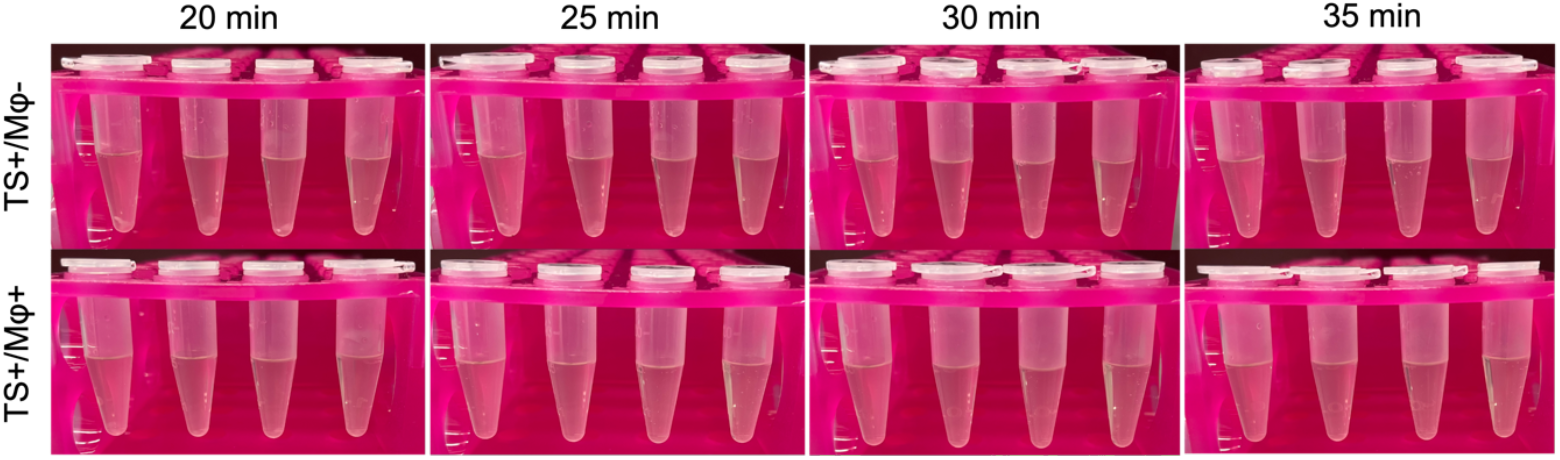
Collagen hydrogel digestion process. Collagen hydrogel was digested in the collagenase solution after 20 min. The gel from the TS^+^/ Mφ^+^ condition was fully digested within 20 min, while the gel from the TS^+^/ Mφ^+^ condition needed more than 30 min to be digested. The four tubes in each photo were four replicates.

**Figure S4.**
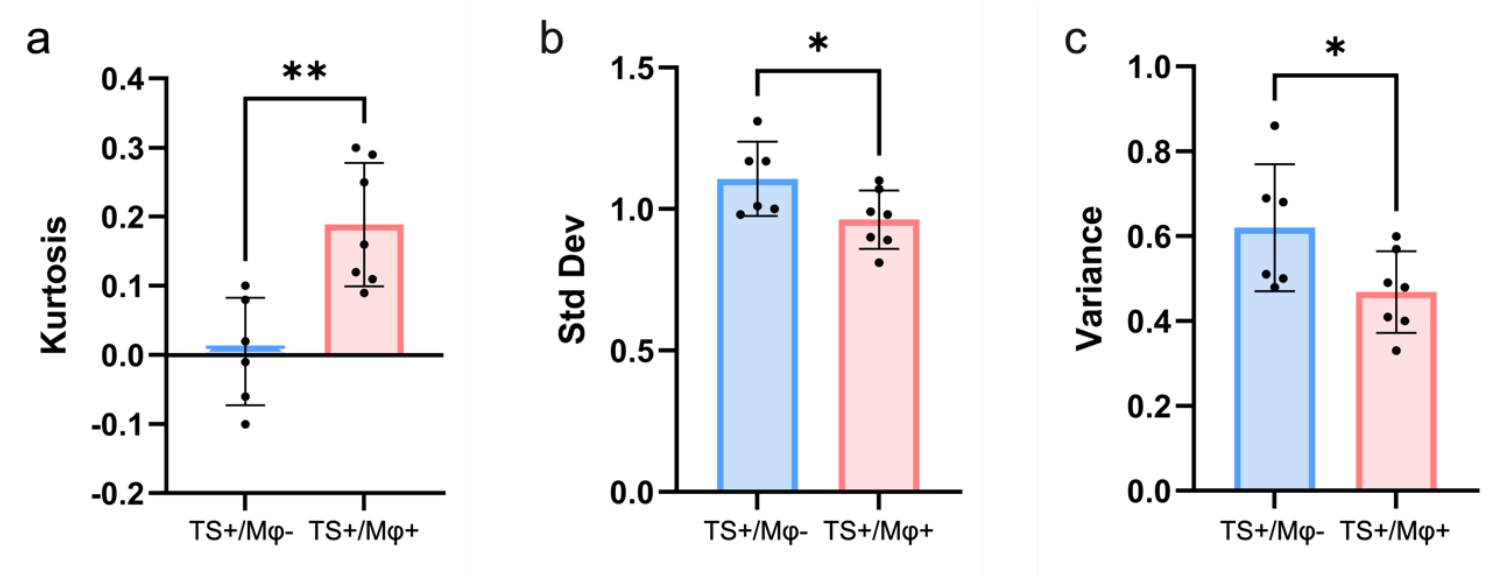
Characterization of collagen fiber orientation. (a) Kurtosis of the fiber orientation distribution. The higher kurtosis indicated a distribution with a sharper peak around the mean and heavier tails, reflecting greater collagen fiber orientation angles concentration near the mean value. (b-c) The lower standard deviation and variance represented the reduced dispersion of collagen fiber orientation angles around the mean, indicating minimal variability within the dataset.

**Figure S5.**
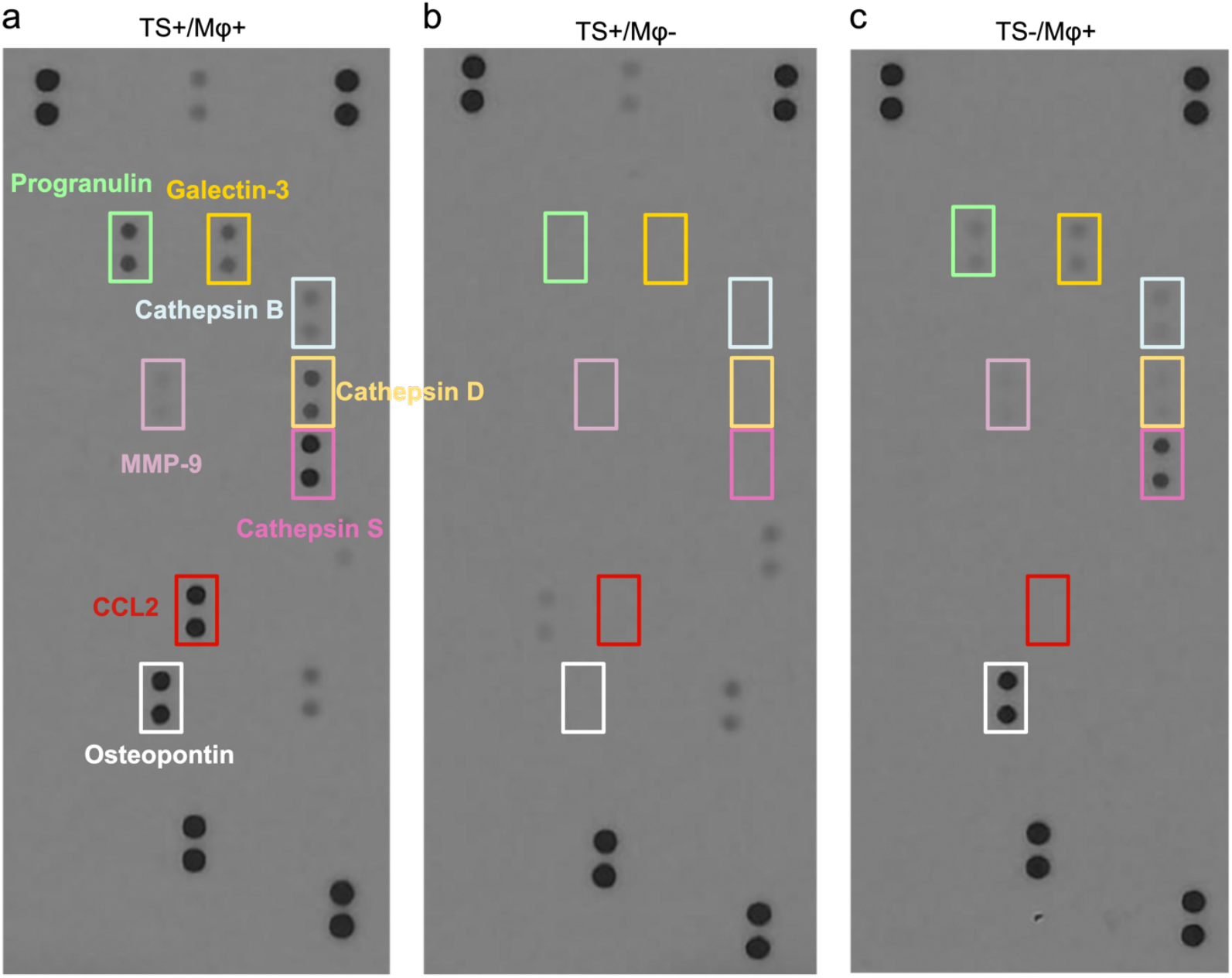
Representative immunoblots from human oncology array. Supernatant was collected from (a) the TS^+^/ Mφ^+^ condition, (b) TS^+^/ Mφ^-^ condition and (c) TS^-^/ Mφ^+^ condition.

## Declaration of Generative AI and AI-assisted technologies in the writing process

During the preparation of this work the author(s) used ChatGPT in order to improve readability and language. After using this tool/service, the author(s) reviewed and edited the content as needed and take(s) full responsibility for the content of the publication.

